# Recruitment of the cellular lipid transport protein CERT to *C. psittaci* inclusions regulates the timing of bacterial egress

**DOI:** 10.1101/2024.11.26.625409

**Authors:** Jana Scholz, Gudrun Holland, Michael Laue, Sebastian Banhart, Dagmar Heuer

## Abstract

Egress of intracellular pathogens is highly regulated and carefully timed. For the zoonotic bacterium *C. psittaci*, the predominant egress pathway is *Chlamydia*-containing sphere (CCS) formation, a calcium-dependent sequential mechanism including protease activity, inclusion membrane destabilization, intracellular calcium increase, and plasma membrane blebbing. How egress is regulated to ensure that it takes place only after *C. psittaci* intracellular development is thus far unknown. Here, we show that *C. psittaci* recruits the cellular ceramide transporter CERT to its inclusion during intracellular development, but this recruitment is reduced at late time points prior to egress. In addition, an early loss of CERT at the inclusion membrane induced by CERT-KO induces premature egress by CCS formation. Complementation of the CERT-KO with different CERT-GFP variants prevents premature egress, except of complementation with a variant lacking the inclusion targeting PH domain, showing that the localization of CERT is critical for CCS formation. The CERT-KO induced premature CCS are formed by the sequential process described for mature CCS, but they contain mostly RBs and are predominantly non-infectious. Thus, our findings suggest that the timing of *C. psittaci* egress by CCS formation is regulated by the recruitment of CERT to the inclusion. We propose that CERT stabilizes the chlamydial inclusion by the formation of ER-inclusion membrane contact sites during intracellular development, and the loss of CERT recruitment facilitates inclusion membrane destabilization and CCS formation.

## Introduction

*Chlamydia psittaci* is a zoonotic pathogen that can be transmitted from birds, which serve as its primary host, to mammalian species, including humans. In humans, *C. psittaci* causes the respiratory disease psittacosis, which is characterized by influenza-like symptoms, including fever, chills, headache, coughing, and dyspnoea. The severity of psittacosis ranges from asymptomatic cases to severe disease and even death.

*C. psittaci* belongs to the genus of *Chlamydia*, a group of Gram-negative, obligate intracellular pathogenic bacteria ^1,2^. During the intracellular development of *Chlamydia*, the bacteria differentiate between two forms: The infectious elementary bodies (EBs) and the non-infectious, replicative reticulate bodies (RBs) ^3–5^. After primary attachment of EBs at the plasma membrane of the host cell, EBs invade by using the host-cellular endocytosis or phagocytosis pathway ^3,6^. Inside of the endocytic vacuole, *Chlamydia* built their replicative niche termed the chlamydial inclusion by type-3 secretion of effector proteins into the inclusion membrane (Inc-proteins) or into the cytosol ^7^. After differentiation of EBs to RBs, rapid replication of *Chlamydia* occurs within the inclusion, and this growth is supported by the acquisition of host cellular metabolites and reorganization of cellular organelles ^8–13^. For example, the host cellular ceramide transporter (CERT) is recruited to *C. psittaci* inclusions and is crucial for inclusion growth and infectious progeny formation ^13^. After rapid replication, RBs redifferentiate to EBs and *C. psittaci* egresses from the host cell, either by lytic egress or by non-lytic *Chlamydia*-containing sphere (CCS) formation ^14^. This sequential process is calcium dependent and involves loss of inclusion membrane integrity, proteolytic activity, an increase in the intracellular calcium concentration, plasma membrane blebbing, and the final detachment of the whole cell as a CCS filled with infectious EBs ^14^. In contrast, other chlamydial species as *C. trachomatis* can perform extrusion formation, another unique non-lytic egress pathway, which is characterized as a cytoskeleton-dependent packaged release process, where both the inclusion membrane and the plasma membrane remain intact and infectious bacteria are released without host cell death ^15–18^.

Besides its functional role as a sphingolipid transporter, CERT has also a mechanistic role during chlamydial infections in stabilizing ER-inclusion membrane contact sites (MCS) ^19,20^. MCS between the ER and chlamydial inclusions were at first characterized for *C. trachomatis* and are supported by the interaction between the bacterial Inc-protein IncD, which is localized in the inclusion membrane and CERT, which localizes to the host ER membrane ^20^. However, no IncD analog of *C. psittaci* is characterized yet, although the recruitment of CERT to *C. psittaci* inclusions was described ^3,13^. For *C. trachomatis*, additional proteins are characterized that stabilize ER-inclusion MCS and thus also stabilize the chlamydial inclusion. These proteins include the ER-resident proteins VAPA/B, interacting with the chlamydial Inc-protein IncV and the ER calcium sensor STIM1, interacting with the chlamydial Inc-protein IncS ^21–28^. Interestingly, STIM1 depletion inhibits egress of *C. trachomatis* by extrusion formation and deletion of IncS of *C. trachomatis* enforces egress by host cell lysis, both suggesting that the stability of ER-inclusion MCS regulate egress of *Chlamydia* spp. ^17,28^.

Here, we examine how CERT recruitment to *C. psittaci* inclusions influences the regulation of *C. psittaci* egress. Our findings demonstrate that the recruitment of CERT to the *C. psittaci* inclusion is diminished in late infections, coinciding with the formation of CCS. The absence of CERT at the inclusion membrane of *C. psittaci* either by knockout of CERT or ectopic expression of a CERT variant lacking the PH domain in CERT-KO cells, resulted in the formation of premature, non-infectious CCS at the mid-infection stage. The findings suggest that the presence of CERT at the inclusion membrane depends on the developmental progress of the bacteria, regulates the stability of chlamydial inclusions, and thereby influences the time point at which egress of *C. psittaci* occurs via CCS formation.

## Results

### CCS formation of *C. psittaci* temporally correlates with a reduction in CERT recruitment

The recruitment of CERT to *C. psittaci* inclusions was exclusively studied for early and middle infections, and not for late infections ^13^. Therefore, this study concentrated on the localization of CERT during late *C. psittaci* infections in the context of bacterial egress by CCS formation. At first, we analyzed CERT recruitment to *C. psittaci* inclusions by performing immunofluorescence staining using antibodies specific for CERT, and bacterial Hsp60. Hsp60-positive *C. psittaci*-inclusions strongly recruit CERT at 24 hours post infection (h pi). However, this recruitment was reduced to weak or no recruitment at 48 h pi (Figure 1 A, right and left inclusion, respectively). Similar to this, ectopically expressed eGFP-CERT localized in close proximity to *C. psittaci* inclusions at 24 h pi, but not at 48 h pi (Figure 1 B).

**Figure 1.**
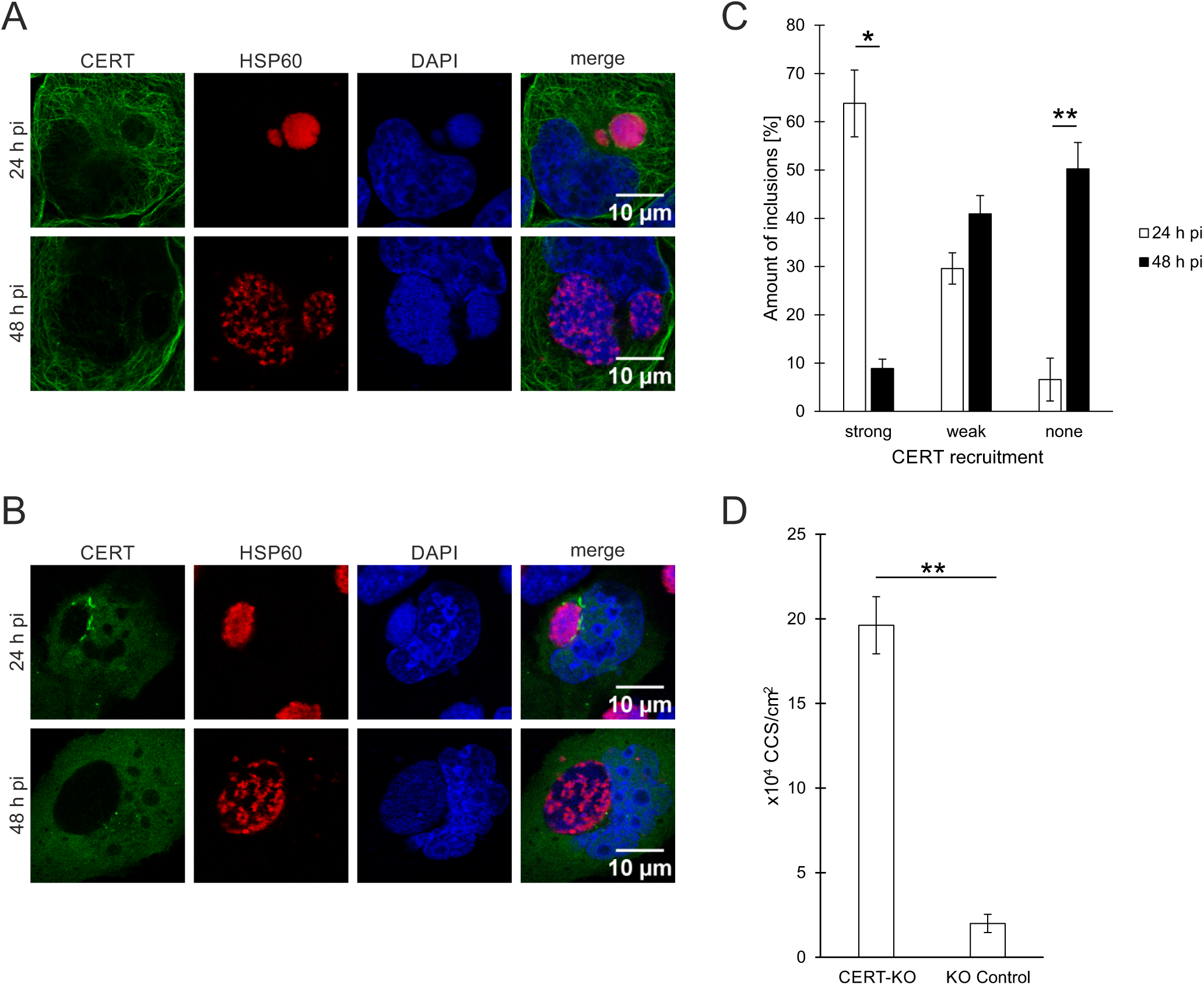
In late infections, CERT recruitment to *C. psittaci* inclusions is reduced and CERT knock out induce early egress of *C. psittaci* by CCS formation at 24 h pi. **(A)** Representative immunofluorescence images of *C. psittaci*-infected HeLa cells (MOI 2) at 24 and 48 h pi. PFA-fixed cells were stained for *C. psittaci* and endogenous CERT using a mouse-anti-Hsp60 (Cy3) and a rabbit-anti-CERT (AF488) antibody, respectively. DNA was counterstained using DAPI. n = 3. **(B)** Representative fluorescence images of HeLa cells transiently expressing eGFP-CERT infected with *C. psittaci* (MOI 2) at 24 and 48 h pi. Cells were PFA-fixed and immunostained for *C. psittaci* using a mouse-anti-Hsp60 (Cy3) antibody and DNA was counterstained using DAPI. n = 3. **(C)** Quantification of CERT recruitment to *C. psittaci* inclusions at 24 and 48 h pi. Based on immunofluorescence images of *C. psittaci*-infected HeLa cells (MOI 2) at 24 and 48 h pi, the number of inclusions showing strong, weak or no CERT recruitment was determined and normalized to the total number of inclusions. At least 60 inclusions per condition were analyzed. Data show mean ± SEM; n = 3; *p < 0.05, **p < 0.01 (Student’s t-test). **(D)** CCS formation in HeLa CERT-KO and KO control cells. Cells were infected with *C. psittaci* (MOI 2), medium was replaced at 20 h pi and CCS in the supernatant were visually quantified at 24 h pi. Data show mean ± SEM; n = 3; **p < 0.01 (Student’s t-test).

Subsequently, the recruitment of CERT was quantified in relation to the inclusions at 24 and 48 hours pi. At 24 h pi, the majority of the inclusions exhibited strong recruitment of CERT (63.8%), 29.6% demonstrated weak CERT recruitment, and only 6.6% were devoid of CERT recruitment (Figure 1C). In contrast, at 48 h pi, the majority of the inclusions did not recruit CERT (50.2%), 40.9% showed a weak CERT-recruitment, and only 8.9% strongly recruited CERT (Figure 1 C).

We therefore sought to determine whether the loss of CERT at the inclusion membrane is associated with egress by CCS formation. To this end, HeLa CERT knockout (KO) cells were infected with *C. psittaci*. We hypothesized that the absence of CERT at the inclusion membrane during intracellular development could lead to premature formation of CCS. We therefore quantified CCS in the supernatant of *C. psittaci*-infected cells at 24 h pi and compared CERT-KO with KO control cells. An almost tenfold increase in formed CCS was observed in the absence of CERT, from 2.0 x 10⁴ CCS/cm² in KO control cells to 19.6 x 10⁴ CCS/cm² in CERT-KO cells at 24 h pi (Figure 1 D).

In conclusion, the data demonstrate that the absence of CERT at the inclusion membrane of *C. psittaci* correlates temporally with egress by CCS formation, indicating that alterations in CERT recruitment to *C. psittaci* inclusions regulate CCS formation.

### Localization of CERT regulates CCS formation

CERT is composed of three functional domains: The N-terminal PH domain, mediating Golgi targeting in uninfected cells, the central FFAT motif, mediating ER targeting, and the C-terminal domain, which is a ceramide-specific lipid-binding START domain (Figure 2 A). During *C. trachomatis*, *C. muridarum*, and *C. psittaci* infections CERT is recruited to the inclusion by its PH-domain ^13,20,29^.

**Figure 2.**
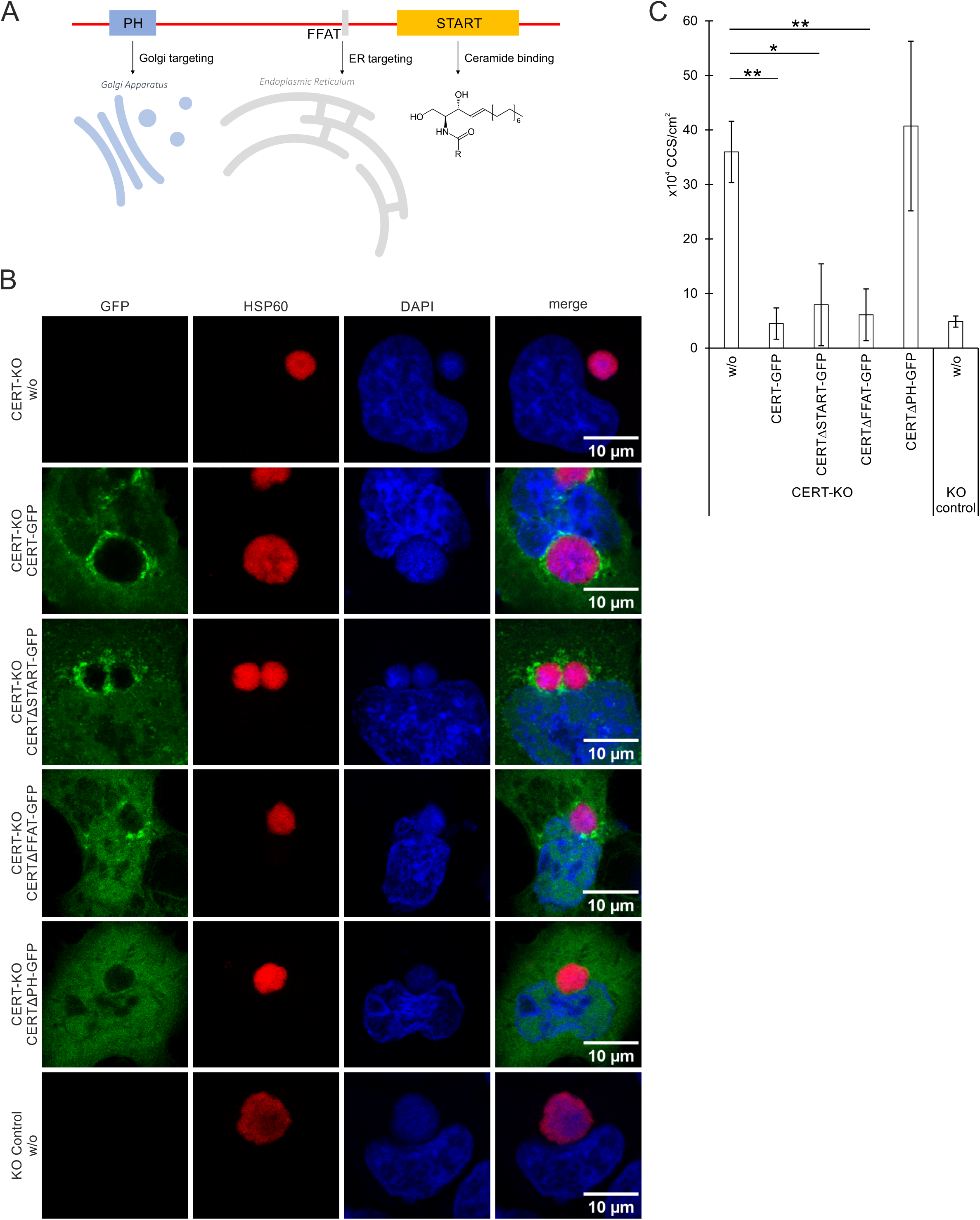
In contrast to full CERT and CERT variants lacking the START- or FFAT-domain, a CERT variant lacking the PH domain is not recruited to *C. psittaci* inclusions and cannot prevent premature CCS formation. **(A)** CERT is composed of the N-terminal PH domain, mediating Golgi targeting in uninfected cells, the central FFAT motif, mediating ER targeting, and the C-terminal START-domain, that is a ceramide-specific lipid binding domain. **(B)** Representative fluorescence images of HeLa CERT-KO cells transiently expressing eGFP-CERT or eGFP-CERT variants lacking the START, FFAT or PH domain and controls infected with *C. psittaci* (MOI 2) at 24 h pi. Cells were PFA-fixed and immunostained for *C. psittaci* using a mouse-anti-Hsp60 (Cy3) antibody; DNA was counterstained using DAPI; n = 4. **(C)** CCS formation in HeLa CERT-KO cells transiently expressing eGFP-CERT or eGFP-CERT variants lacking the START, FFAT or PH domain and controls. Cells were infected with *C. psittaci* (MOI 2), medium was replaced at 20 h pi and CCS in the supernatant were visually quantified at 24 h pi. CCS number was normalized on the transfection efficiency. Data show mean ± SEM; n = 4; *p < 0.05, **p < 0.01 (Student’s t-test).

Initially, we infected CERT-KO cells ectopically expressing distinct eGFP-CERT variants with *C. psittaci* and examined the localization of the eGFP-CERT variants through immunofluorescence staining at 24 h pi. In cells that had not been transfected with the CERT-KO construct, no GFP signal was observed (Figure 2B). In CERT-KO cells transfected with full-length eGFP-CERT, the protein was observed to localize in close proximity to *C. psittaci* inclusions (Figure 4B), a finding that is comparable to those observed in wildtype HeLa cells (Figure 1B). A comparable staining pattern was observed following transfection with eGFP-CERT lacking the START or FFAT domain (Figure 2B). As published, eGFP-CERT lacking the PH domain was not recruited to *C. psittaci* inclusions ^13^.

Next, we analyzed CCS formation after infecting CERT-KO cells ectopically expressing different eGFP-CERT variants with *C. psittaci*. In untransfected CERT-KO cells, 36.0 x 10^4^ CCS/cm^2^ were formed compared to 4.9 x 10^4^ CCS/cm^2^ in untransfected KO control cells (Figure 2 C). In the conditions in which the CERT variants were introduced by transient transfection, the CCS numbers were normalized to the transfection efficiencies. Upon transfection of CERT-KO cells with full length eGFP-CERT, the normalized CCS formation (4.5 x 10^4^ CCS/cm^2^) resembles that detected in the control cells, indicating that full length CERT can complement the premature egress phenotype seen in the CERT-KO cells. Similarly, transfection of CERT-KO cells with eGFP-CERT lacking the START or FFAT domain significantly decreased CCS formation to 7.9 x 10^4^ and 6.1 x 10^4^ CCS/cm^2^, respectively. In contrast, transfection of CERT-KO cells with eGFP-CERT lacking the PH domain did not decrease CCS formation (40.7 x 10^4^ CCS/cm^2^) (Figure 2 C).

Taken together, expression of CERT variants revealed that the ability of CERT to be recruited to the inclusion correlates with its capacity to prevent premature egress and suggest that CERT recruitment to the inclusion controls CCS formation.

### CERT-KO induces early CCS formation by a sequence of events analogous to mature CCS formation

Next, we asked if the early egress seen in infected CERT-KO cells has the same mechanistic characteristics as the (mature) CCS formation that occurs in wildtype cells at the culmination of *C. psittaci* development. Mature CCS are formed by a process with distinct characteristics ^14^. An increase in intracellular calcium precedes protease activity that can be detected by cleavage of a DEVD tetrapeptide-containing substrate. Loss of inclusion membrane integrity can be detected by recruitment of EqtSM caused by exposure of sphingomyelin to the cytoplasm. Plasma membrane blebbing then leads to eventual detachment of the CCS^14^. Thus, we examined the sequence of events described for mature CCS during the early CCS formation induced in CERT-KO cells.

At first, we monitored early CCS formation using HeLa CERT-KO cells transiently expressing eGFP starting at 20 h pi. Before CCS formation, the bacterial inclusion excluded the cytosolic eGFP (Figure 3 A). Then in all analyzed cells, CCS formation initiated with blebbing of the cellular plasma membrane and influx of eGFP into the inclusion lumen. This was followed by enlargement of the plasma membrane blebs and subsequent detachment of the entire host cell, which completed the formation of the CCS in the supernatant of the cell culture.

**Figure 3.**
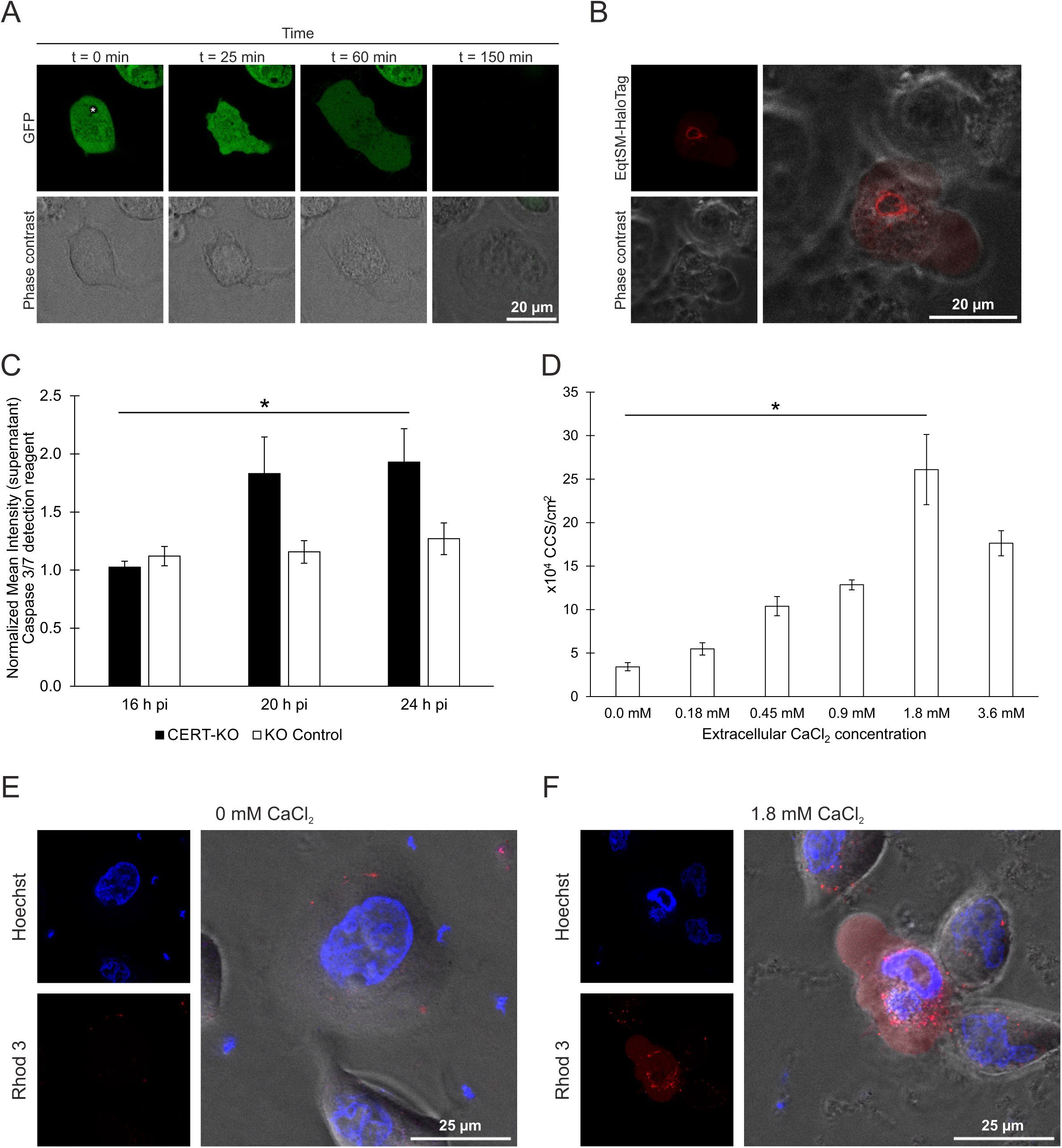
CERT-KO-induced early CCS are formed by an analogous sequence of events as mature CCS. **(A)** During early CCS formation, plasma membrane blebbing occurs and the inclusion membrane destabilizes. HeLa CERT-KO cells transiently expressing eGFP were infected with *C. psittaci* (MOI 2) and monitored live starting at 20 h pi using a CLSM. Panels show representative images of a section plane of a CCS-forming CERT-KO cell; The inclusion is marked with an asterisk; n = 3. **(B)** EqtSM is recruited to *C. psittaci* inclusions during early CCS formation. CERT-KO cells transiently expressing EqtSM-HaloTag were infected with *C. psittaci* (MOI 2). At 20 h pi, cells were labeled with 200 nM Janelia Fluor 585 HaloTag Ligand (Promega) and monitored live. Panel shows representative images of an EqtSM-HaloTag-recruiting *C. psittaci* inclusion during early CCS formation; n = 3. **(C)** A DEVD-cleaving protease is activated during early CCS formation. *C. psittaci*-infected CERT-KO and KO control cells (MOI 2) were stained with Incucyte Caspase-3/7 Dye for Apoptosis at 16 h pi and monitored using a CLSM for 8 h. Mean fluorescence intensity of infected cells at 16, 20, and 24 h pi was normalized to mean fluorescence intensity of uninfected cells at respective time points. Data show mean ± SEM; n = 3; *p < 0.05 (Student’s t-test). **(D)** Early CCS formation depends on the extracellular calcium concentration. CERT-KO cells were infected with *C. psittaci* (MOI 2). At 20 h pi, culture medium was replaced by serum-free medium supplemented with the indicated concentrations of calcium chloride. CCS in the supernatant were visually quantified at 24 h pi. Data show mean ± SEM; n = 3; *p < 0.05 (Student’s t-test). **(E, F)** CERT-KO cells were infected with *C. psittaci* (MOI 2). At 20 h pi, medium was replaced by serum-free medium supplemented with 0 mM calcium chloride **(E)** or 1.8 mM calcium chloride **(F)**. At 22 h pi, cells were labeled with the calcium sensor Rhod-3 for 1 h. At 24 h pi, DNA was counterstained using Hoechst and images were acquired. Representative images of intact cells (0.0 mM CaCl_2_) and CCS formation (1.8 mM CaCl_2_) are shown; n = 2.

Next, we asked if EqtSM is recruited to the *C. psittaci*-inclusion when early CCS formation starts. To test this, we transiently expressed EqtSM-HaloTag in CERT-KO cells, labeled them with HaloTag Ligand and performed live cell imaging starting at 20 h pi. We observed many cells recruiting EqtSM to the inclusion coincident with the start of plasma membrane blebbing (Figure 3 B) and followed by CCS formation.

In addition, we tested if proteolytic cleavage of a DEVD-containing substrate is increased in adherent CERT-KO cells during early CCS formation. Indeed, we detected a significant increase in fluorescence intensity of a cleaved DEVD-containing substrate compared to uninfected cells from 1.0-fold at 16 h pi to 1.8-fold and 1.9-fold at 20 and 24 h pi, respectively (Figure 3 C). In comparison, in control cells the intensity was only slightly increased from 1.1-fold at 16 h pi to 1.2-fold and 1.3-fold at 20 and 24 h pi, respectively.

Furthermore, the role of calcium signaling for early CCS formation was investigated. During mature CCS formation, CCS numbers increased with extracellular calcium concentrations ^14^. Similarly, during early CCS formation in CERT-KO cells, we observed a dose-dependent increase in CCS numbers from 3.4 x 10^4^ CCS/cm^2^ to 26.1 x 10^4^ CCS/cm^2^ when increasing the extracellular calcium from 0.0 to 1.8 mM CaCl_2_ (Figure 3 D).

Subsequently, we analyzed the intracellular calcium concentration during early CCS formation using the Rhod-3 membrane-permeant cytosolic calcium sensor under conditions containing 0.0 and 1.8 mM calcium chloride. At 0.0 mM calcium chloride, we did not detect an increase in cytosolic calcium concentration in *C. psittaci*-infected CERT-KO cells at 24 h pi (Figure 3 E). However, at 1.8 mM calcium chloride, we observed *C. psittaci*-infected CERT-KO cells with blebbing membranes and an increased cytosolic calcium concentration (Figure 3 F).

In sum, this data revealed that CERT-KO induced early CCS are formed by an analogous sequence of events as mature CCS including plasma membrane blebbing, inclusion membrane destabilization, EqtSM recruitment, activation of a DEVD cleaving protease, dependency on the extracellular calcium concentration and intracellular calcium increase. This further supports that the presence and absence of CERT regulates CCS formation.

### CERT-KO induces the formation of RB-containing, non-infectious CCS

At 24 h pi, *C. psittaci* inclusions contain mostly RBs, with only few bacteria starting to redifferentiate to EBs ^30^. Thus, we asked if bacteria in early-formed CCS are infectious. To test this, we separated CCS from free bacteria present in the supernatant of *C. psittaci*-infected CERT-KO and control cells at 24 h pi using differential centrifugation. We then quantified the infectious progeny (IFU) relative to the bacterial genome copy numbers associated with the harvested CCS. Although more CCS are released in the supernatants from CERT-KO cells than from control cells 24 h pi, the percentage of the progeny that are infectious are comparable (0.003 ± 0.0016% and 0.006 ± 0.0038%, respectively) (Figure 4 A). These values 24 h pi are extremely reduced compared to the percentage of infectious progeny in mature CCS released from HeLa cells at 48 h pi (3.2 ± 0.9%) ^14^.

**Figure 4.**
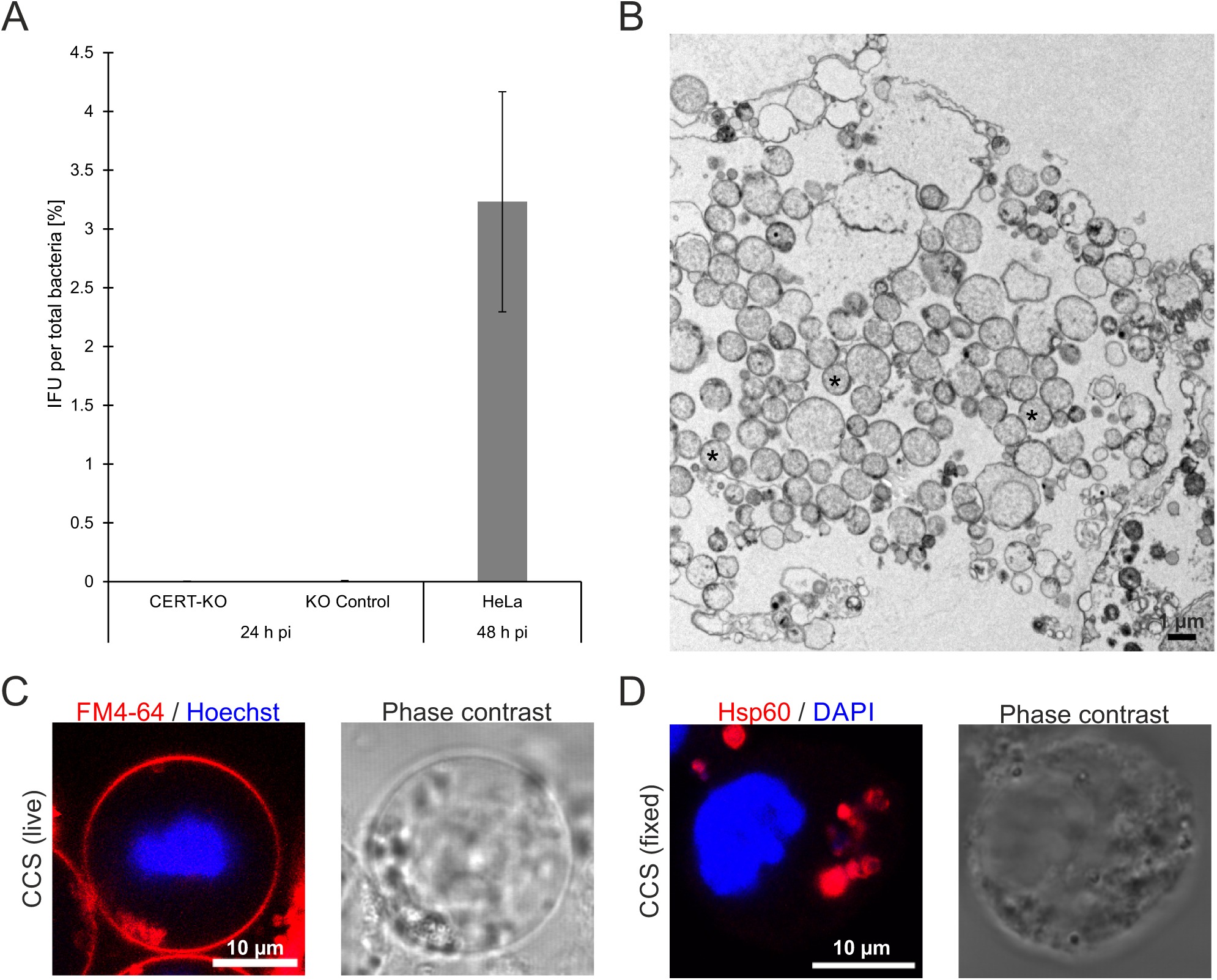
Premature, CERT-KO-induced CCS contain mainly non-infectious reticulate bodies. **(A)** CCS in the supernatant of *C. psittaci*-infected CERT-KO and KO control cells (MOI 2) were separated from free bacteria by centrifugation (300 x *g*, 5 min, RT) at 24 h pi. Infectious progeny was titrated after glass bead lysis and numbers were normalized to genome copy numbers determined by qRT-PCR. Data show mean ± SEM; n = 3. **(B)** Transmission electron microscopy (TEM) of a thin section through the chemically fixed supernatant of *C. psittaci*-infected HeLa CERT-KO cells (MOI 2, 24 h pi). Numerous *Chlamydia*, mostly in RB stage, were found; n = 3. The image shows a group of RBs (*) which are associated with cellular debris, which is the main component of the supernatant. **(C)** Representative fluorescence images of an early live CCS isolated of the supernatant of *C. psittaci*-infected HeLa CERT-KO cells (MOI 2, 24 h pi). The surrounding membrane was visualized using the membrane marker FM 4-64 and DNA was counterstained by Hoechst; n = 3. **(D)** Representative fluorescence images of a PFA-fixed early CCS isolated from *C. psittaci*-infected HeLa CERT-KO cells (MOI 2, 24 h pi). Bacteria inside the CCS were detected using a chlamydial Hsp60 (Cy3) antibody and the DNA was counterstained using DAPI; n > 3.

To assess the presence and morphology of the *C. psittaci* released from infected CERT-KO cells, we analyzed the culture supernatants using thin-section transmission electron microscopy (TEM). Among cellular debris, which is the main component of the supernatant, we found numerous *Chlamydia*. which were, with few exceptions, in RB stage (Figure 4 B). We further analyzed the morphology of early CCS by staining with the membrane dye FM 4-64 and with Hoechst. Fluorescence live cell confocal microscopy revealed that early CCS were surrounded by a membrane and contained concentrated DNA, which are both features of mature CCS (Figure 4 C) ^14^. Bacteria could be detected in fixed early CCS with antibodies specific for bacterial Hsp60 (Figure 4 D). Bacteria were also labeled with the DNA marker DAPI, which also labeled DNA we identified as the host cell nucleus, which is also found within mature CCS ^14^.

These data show that CCS released early in the absence of CERT rarely contain any infectious EBs although the CCS are morphologically similar to mature CCS. In wildtype infections, CERT recruitment is reduced after RB redifferentiation to the EB stage, indicating that stage-controlled changes of the localization of CERT allow release of infectious progeny containing CCS.

## Discussion

The stability of pathogen-containing vacuoles is crucial for intracellular vacuolar pathogens. On the one hand, stabilization of the vacuole facilitates pathogen growth in a protected niche: on the other hand, the vacuole must allow pathogen egress from host cells and transmission to other hosts ^31–33^. For the zoonotic bacterial pathogen *C. psittaci*, the predominant egress strategy is via CCS formation, a process where bacteria are released from a destabilized inclusion to the host cytosol and the whole host cell subsequently detaches from the remaining cellular monolayer as a CCS ^14^. This process depends on the extracellular calcium concentration, bacterial protease activity and a rapid increase in the cytosolic calcium concentration ^14^. However, it remains unclear how the stage-dependent stabilization and destabilization of the inclusion membrane is regulated to ensure completion of intracellular development before egress by CCS formation.

In this study, we showed that CERT recruitment to the inclusion membrane is time-dependent: During RB replication at 24 h pi, CERT is recruited to the inclusion membrane. Here it can stabilize ER-inclusion MCS ^19,20^. In contrast, during CCS formation at 48 h pi, CERT recruitment is reduced and the inclusion loses stability (Figure 5). Premature loss of CERT recruitment, which can be induced by using CERT-KO cells, induces the formation of premature, non-infectious RB-containing CCS by a sequence of events analogous to mature CCS formation (Figure 5) ^14^. We propose that the time-dependent recruitment of CERT stabilizes the inclusion by ER-inclusion MCS during RB replication and that reduction of CERT at the inclusion after EB maturation allows destabilization of the inclusion by the loss of ER-inclusion MCS prior to CCS formation.

**Figure 5.**
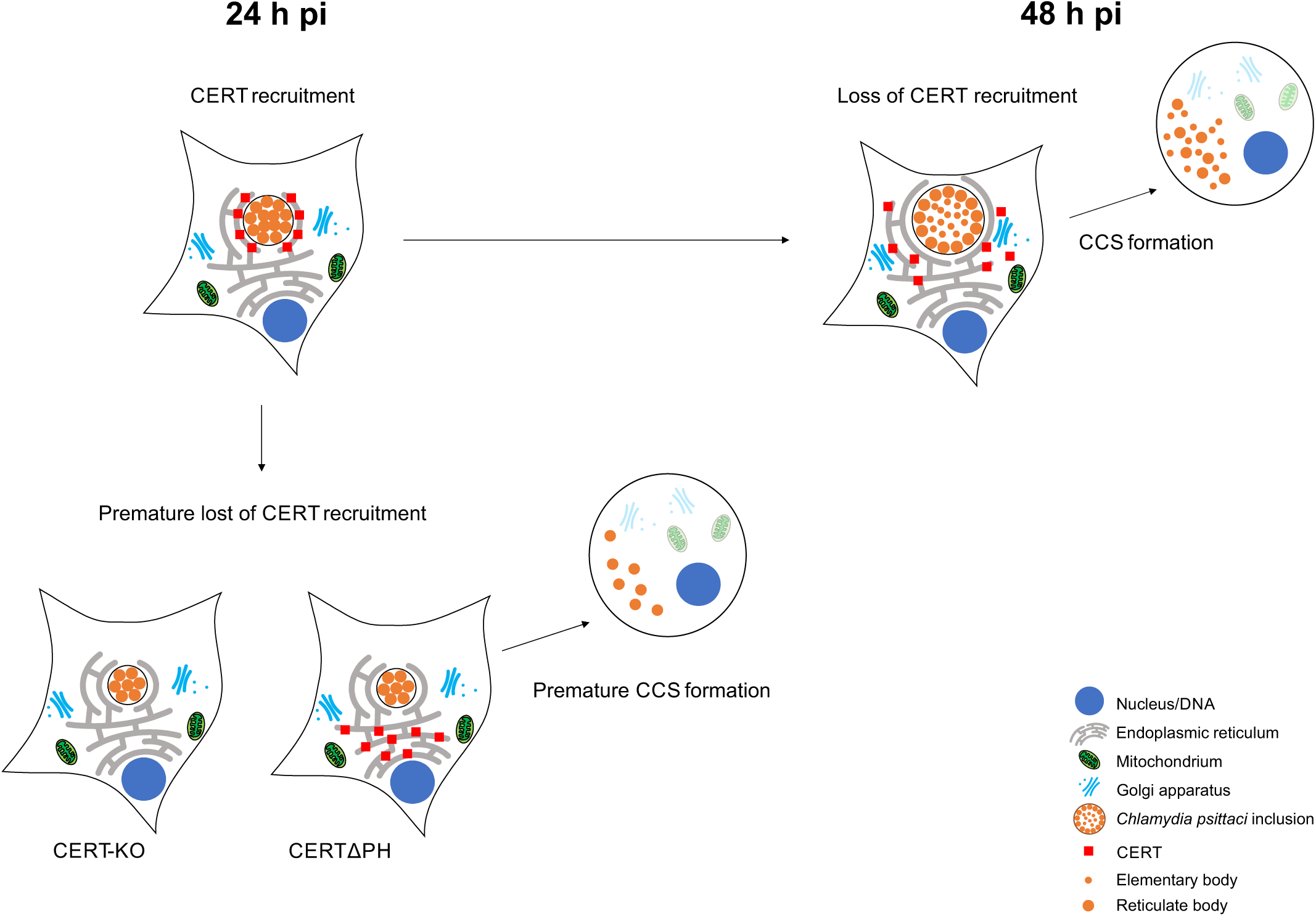
Graphical model of the role of CERT in regulating *C. psittaci* CCS formation. At 24 h pi, the recruitment of CERT to the *C. psittaci* inclusion promotes inclusion stability by CERT-stabilized ER-inclusion membrane contact sites. At 48 h pi, CERT recruitment to the *C. psittaci* inclusion is reduced and the consequential loss of the inclusion stability facilitates CCS formation. When CERT is not present at the inclusion membrane at 24 h pi (CERT-KO or CERTΔPH), this promotes early egress and facilitates the formation of premature, non-infectious CCS.

The role stability of ER-inclusion MCS plays in egress of *Chlamydia* spp. was previously discussed in terms of the IncS-STIM1 interaction at *C. trachomatis* inclusions ^25,27,28^. *C. trachomatis* can egress from infected cells via extrusions, and unlike during CCS formation, the inclusion membrane remains intact when extrusions form. Destabilization of ER-inclusion MCS either by a lack of STIM1 induced by siRNA-mediated knock-down or by a lack of *C. trachomatis* IncS shifted *C. trachomatis* egress to reduce extrusion formation or increase inclusion lysis, respectively ^17,28^. Notably, *C. psittaci* does not recruit STIM1 at 24 and 48 h pi (Figure S1) and extrusion formation takes not place in *C. psittaci* infections ^14^. Perhaps the absence of STIM1-containing MCS result in reduced stability of the *C. psittaci* inclusions compared to *C. trachomatis* inclusions, and this feature could preclude *C. psittaci* extrusion formation. Following this model, CERT recruitment would be more important for the stabilization of *C. psittaci* ER-inclusion MCS compared to *C. trachomatis* ER-inclusion MCS, which may explain the influence CERT recruitment has in regulating CCS formation in *C. psittaci* in a time-dependent manner. Interestingly, STIM1 is also not recruited to *C. muridarum* inclusions ^28^. It is therefore tempting to speculate that also in *C. muridarum* egress could be regulated by the recruitment of CERT.

CERT does not only stabilize ER-inclusion MCS in chlamydial infections, but it also plays a crucial role for sphingolipid acquisition of *Chlamydia* spp ^13,20,29,34–36^. In *C. trachomatis*, *C. muridarum* and *C. psittaci* infections, chemical inhibition of CERT with the ceramide analogue HPA-12 reduces inclusion size and progeny formation ^13,20,29^. Interestingly, CERT-KO does not prevent sphingolipid acquisition at the inclusion in *C. psittaci* infections, but HPA-12 treatment does, indicating that a CERT-independent, HPA-12-sensitive route of ceramide acquisition is used by *C. psittaci* in CERT-KO cells ^13^. Accordingly, EqtSM recruitment to sphingomyelin exposed to the inclusion membrane prior to CCS formation can be observed during CCS formation in CERT-KO cells. In addition, expression of a CERT variant lacking the START domain, which is not able to transport ceramides, but localizes at the inclusion membrane, can compensate for the CERT-KO-induced premature CCS formation. These results further indicate that premature CCS formation in the absence of CERT is due to the loss of its mechanistic role in stabilizing ER-inclusion MCS and is not based on its role in sphingolipid acquisition during *C. psittaci* infection.

To further understand how the time-dependent recruitment of CERT is regulated, it will be crucial to identify the bacterial interaction partner of CERT within the *C. psittaci* inclusion membrane. For several chlamydial species, including *C. trachomatis*, *C. muridarum*, *C. suis*, *C. caviae*, and *C. felis*, the bacterial Inc-protein IncD interacts with the PH domain of CERT at the inclusion membrane ^13,21,29,34^. However, no IncD analog has yet to be identified in *C. psittaci*, even though the PH domain of CERT is involved in its recruitment to the *C. psittaci* inclusion and in its interaction with IncD ^3,13^.

Taken together, we showed that the recruitment of CERT stabilizes *C. psittaci* inclusions during RB replication and prevents premature CCS formation. Our data suggest that loss of CERT recruitment in late infections facilitates the destabilization of ER-inclusion MCS and the chlamydial inclusion membrane and consequently promotes CCS formation and bacterial egress.

## Material and Methods

### Cell culture, transient transfection and infection assays

Cell culture, transient transfection and infection assays were performed as described previously ^13,14^. Briefly, HeLa cells (ATCC CCL-2) were cultivated in RPMI 1640 medium (Gibco) supplemented with 10% fetal bovine serum (FBS, Sigma-Aldrich), 1 mM sodium pyruvate (Gibco), and 2 mM L-glutamine (Gibco) at 37°C and 5% CO_2_ and passaged every 2 to 3 days. Transient transfection was performed with Lipofectamine 2000 reagent (Thermo Fisher) 4 h prior to infection according to manufacturer’s instructions. For infection assays, subconfluent HeLa cells were washed with infection medium (Dulbecco’s modified Eagle’s medium (DMEM, Gibco) with glucose (4.5 g L^-1^) supplemented with 5% FBS, 1 mM sodium pyruvate, and 2 mM L-glutamine) and incubated at 35°C and 5% CO_2_ using the indicated multiplicity of infection (MOI) with *C. psittaci* strain 02DC15 ^37^. *C. psittaci*-infected cell cultures were centrifuged (30 min, 600 x *g*, room temperature (RT)) at 30 min pi and washed with infection medium at 2 h pi.

### Quantification of chlamydial genome copy number (GCN) by quantitative real-time PCR

Determination of bacterial genome copy numbers by quantitative real-time PCR (qRT-PCR) was performed as already described ^38^.

### Quantification of CCS

The number of CCS in *C. psittaci* infected cell cultures was determined as described previously ^14^. In brief, cell cultures were washed with PBS at indicated time points and medium was changed to conditional medium and cultivation was continued for additional 4 h. The number of CCS in the supernatant of the infected cell cultures was quantified in 12 well plates. The average number of CCS of at least five visual fields was determined and normalized to the infected cell area.

### Live cell imaging of premature *C. psittaci* egress in CERT-KO cells

To monitor early CERT-KO-induced egress of *C. psittaci*, live cell imaging of *Chlamydia*-infected CERT-KO cells transiently expressing eGFP was performed as described previously ^14^. In brief, live cell microscopy of infected cells cultured in 8 well chambered coverslips (µ-Slide 8 Well, Ibidi) was performed at a Stellaris 8 Confocal Microscope (Leica Microsystems) equipped with a live cell chamber at 35°C. Z-stacks of 12 slices with each 3 µm distance covering both adherent cells and the supernatant containing CCS were acquired every 2-2.5 minutes from 20 to 24 h pi.

### Imaging of cytosolic sphingomyelin (SM) exposure

To image the cytosolic SM exposure at the inclusion membrane, CERT-KO cells transiently expressing a non-toxic, HaloTagged version of equinatoxin II (EqtSM-HaloTag) as cytosolic SM reporter were used ^40,41^. Cell culture, transient transfection and infection with *C. psittaci* (MOI 2) were performed in 8 well chambered coverslips (µ-Slide 8 Well, Ibidi). At 19.5 h pi, cells were labeled with 200 nM Janelia Fluor 585 HaloTag Ligand (Promega) in infection medium for 30 min. Subsequently, cells were washed with infection medium and live cell imaging was performed at a Stellaris 8 Confocal Microscope (Leica Microsystems) equipped with a live cell chamber at 35°C. Z-stacks of 12 slices with each 3 µm distance were acquired every 2:10 minutes for a total of 4 hours.

### Determination of proteolytic DEVD cleaving activity

Determination of proteolytic DEVD cleaving activity was performed as described previously ^14^. For this, the Incucyte Caspase-3/7 Red Dye for Apoptosis (Sartorius) was used as described in the manufacturer’s protocol. Cells were cultured and infected with *C. psittaci* (MOI 2) in 8 well chambered coverslips (µ-Slide 8 Well, Ibidi). Incucyte Caspase-3/7 Red Dye was diluted in infection medium to a final assay concentration of 833 nM and added to *C. psittaci*-infected cell cultures at 16 h pi. Live imaging was performed using an LSM 780 CLSM (Carl Zeiss) equipped with a live cell chamber at 35°C. Z-stacks of 12 slices with each 3 µm distance covering both adherent cells and CCS were acquired every 5 minutes for a total of 8 hours.

### Calcium imaging

Imaging of intracellular calcium levels and distribution was performed by Rhod 3 staining using the Rhod-3 Calcium Imaging Kit (Thermo Fisher Scientific) following the manufacturer’s protocol as described previously ^14^. In brief, CERT-KO cells cultured in 8 well chambered coverslips (µ-Slide 8 Well, Ibidi) were infected with *C. psittaci* (MOI 2) and at 20 h pi, infected cells were washed with PBS and medium was changed to calcium- and FBS-free infection medium (Cat. No. 21068028, Gibco) supplemented with the indicated concentrations of calcium chloride (Roth). At 22 h pi, cells were washed with PBS twice and 150 µL per well of freshly prepared Rhod-3 loading buffer (1x PowerLoad concentrate, 10 µM Rhod-3 AM, 250 µM Probenecid in infection medium or calcium- and FBS-free infection medium) were added. After incubation (60 min, 35°C, 5% CO_2_), cells were washed three times with PBS and 150 µL per well of incubation buffer (250 µM Probenecid in infection medium or calcium- and FBS-free infection medium) were added. After incubation (60 min, 35°C, 5% CO_2_), cells were washed with PBS and 150 µL per well of 25 mg mL^-1^ Hoechst 33342 (Sigma-Aldrich) diluted 1:2000 in infection medium were added for live cell microscopy. Live cell microscopy was performed at a Stellaris 8 Confocal Microscope (Leica Microsystems).

### Separation of CCS and free bacteria for reinfection assay

HeLa CERT-KO and KO control cells were initially infected with *C. psittaci* and incubated for 24 h. Then, supernatants were collected and centrifuged at low speed (5 min, 300 x *g*, RT) to pellet CCS and separate them from free bacteria in the remaining supernatant. Subsequently, CCS and free bacteria were subjected to glass bead lysis followed by infection of freshly seeded HeLa cells. Cells were fixed at 24 h post infection (pi), stained for Hsp60 (see immunofluorescence assay for more details), and numbers of infection-forming units (IFU) per mL were calculated from average inclusion counts in 10 fields of view per condition. IFU were normalized on GCN (see quantification of chlamydial genome copy number (GCN) by quantitative real-time PCR for more details).

### Transmission electron microscopy

Transmission electron microscopy was performed as described previously ^14^. In brief, at 24 h pi the culture supernatant of *C. psittaci*-infected (MOI 2) HeLa CERT-KO cells (sedimented by centrifugation for 5 min at 300 x *g*) were fixed with a mixture of glutaraldehyde and formaldehyde (2.5% and 1%, respectively; in 50 mM HEPES buffer). After incubation for 2 h at RT, suspensions were centrifuged (6000 x *g*, 10 min) and sediments were embedded in low-melting point agarose. Small sample blocks were extracted from the solidified agarose, post fixed in osmium tetroxide (1% in distilled water) followed by block contrasting with tannic acid (0.1% in 50 mM HEPES buffer) and uranyl acetate (2% in distilled water). Subsequently, samples were dehydrated by incubation in a graded ethanol series and embedded in epon resin. After polymerization for 48 h at 60 °C, ultrathin sections (approx. 70 nm) were prepared using an ultramicrotome (UC7; Leica) and stained with uranyl acetate (2% in distilled water, 20 min) followed by lead citrate (2 min) to increase contrast. Sections were examined with a transmission electron microscope (Tecnai 12, FEI Corp.) at 120 kV. Images were recorded using a CMOS-camera (Phurona, Emsis).

### Isolation and staining of CCS

CCS in the supernatant of *C. psittaci*-infected cell cultures were separated by centrifugation (5 min, 300 x *g*, RT) at 24 h pi. For live staining, pellet was mixed with 10 mM FM4-64 in DMSO (Sigma-Aldrich) diluted 1:1000 in infection medium and 25 mg mL^-1^ Hoechst 33342 (Sigma-Aldrich) diluted 1:2000 in infection medium. CCS were transferred into a 15 well chambered coverslip (µ-Slide 15 Well 3D, Ibidi) and examined using an LSM 780 CLSM (Carl Zeiss). For immunofluorescence staining, pellet was mixed with 4% paraformaldehyde in PBS and transferred into a poly-L-lysine coated 8 well chambered coverslip (µ-Slide 8 Well, Ibidi). After 30 min of incubation at RT, staining was continued as described for immunofluorescence assays.

### Immunofluorescence assay

For immunofluorescence assays, *C. psittaci*-infected (MOI 2) or uninfected HeLa cells, HeLa CERT-KO cells or HeLa KO control cells were fixed with 4% paraformaldehyde in PBS (30 min, RT) at the indicated time points. After washing with PBS, samples were blocked and permeabilized using 0.2% Triton X-100 and 0.2% BSA in PBS (20 min, RT). Incubation with primary antibodies (1 h, RT) was followed by washing and incubation with secondary antibodies and DAPI (25 µg ml^-1^, Sigma-Aldrich) (1 h, RT). All antibodies were diluted using 0.2% BSA in PBS. Antibodies were used at the following concentrations: mouse anti-Hsp60 (chlamydial, cHsp60; 1:600, Cat. No. ALX-804-072, Enzo Life Sciences), rabbit anti-CERT (1:200, Cat. No. PA5-115035, Invitrogen), rabbit anti-STIM1 (1:50, Cat. No. 11565-1-AP, Proteintech), Alexa Fluor 488-coupled goat anti-rabbit IgG (1:100, Cat. No. 111-545-144, Dianova), and Cy3-coupled goat anti-mouse IgG (1:200, Cat. No. 115-165-146, Dianova). All samples were analyzed at a Stellaris 8 Confocal Microscope (Leica Microsystems).

## Acknowledgements

We thank Andrea Martini, Madita Winkler and Henning Krüger for excellent technical assistance. The EqtSM-HaloTag expression vector as cytosolic SM reporter was kindly provided by Michael Hensel (Universität Osnabrück). This work was financially supported by the Deutsche Forschungsgemeinschaft (www.dfg.de): SPP 2225 to Dagmar Heuer (HE 6008/4-1).

## Author contributions

D.H. conceptualized the study. J.S., E.R., and S.B. performed the experiments. G.H. and M.L. performed the EM studies. J.S., D.H., S.B., M.L. and G.H. analyzed the data. J.S. and D.H. wrote the original manuscript. All authors reviewed the manuscript and approved the final version.

## Data availability statement

The datasets generated during and/or analysed during the current study are available from the corresponding author on reasonable request.

---

**Supplemental Figure S1:**
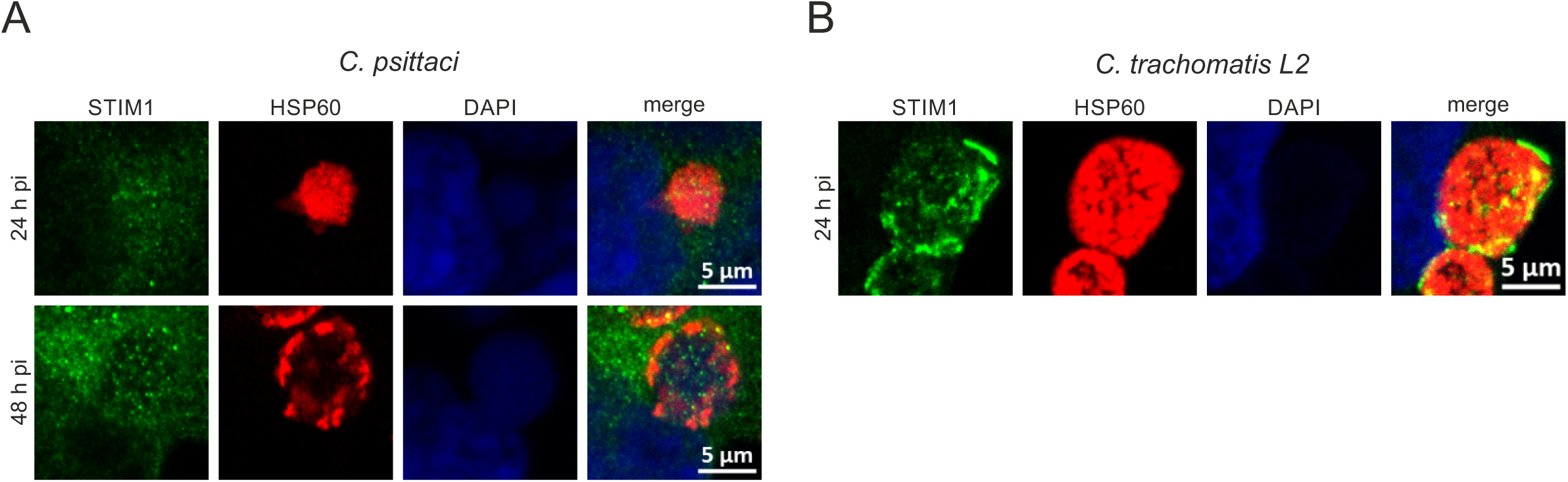
In contrast to *C. trachomatis* L2, *C. psittaci* does not recruit STIM1 to the chlamydial inclusion. **(A)** Representative immunofluorescence images of *C. psittaci*-infected HeLa cells (MOI 2) at 24 and 48 h pi. PFA-fixed cells were stained for *C. psittaci* and endogenous STIM1 using a mouse-anti-Hsp60 (Cy3) and a rabbit-anti-STIM1 (AF488) antibody, respectively. DNA was counterstained using DAPI. n = 3. (B) Representative immunofluorescence images of *C. trachomatis* L2-infected HeLa cells (MOI 2) at 24 h pi. PFA-fixed cells were stained for *C. trachomatis* L2 and endogenous STIM1 using a mouse-anti-Hsp60 (Cy3) and a rabbit-anti-STIM1 (AF488) antibody, respectively. DNA was counterstained using DAPI. n = 3.

## References

1. Bachmann, N. L., Polkinghorne, A. & Timms, P. *Chlamydia* genomics: providing novel insights into chlamydial biology. Trends Microbiol. 22, 464–472 (2014).

2. Bayramova, F., Jacquier, N. & Greub, G. Insight in the biology of *Chlamydia*-related bacteria. Microbes Infect. 20, 432–440 (2018).

3. Banhart, S., Schäfer, E. K., Gensch, J. M. & Heuer, D. Sphingolipid Metabolism and Transport in *Chlamydia trachomatis* and *Chlamydia psittaci* Infections. Front. Cell Dev. Biol. 7, 1–8 (2019).

4. Knittler, M. R. & Sachse, K. *Chlamydia psittaci*: Update on an underestimated zoonotic agent. Pathog. Dis. 73, 1–15 (2015).

5. Radomski, N., Einenkel, R., Müller, A. & Knittler, M. R. *Chlamydia*-host cell interaction not only from a bird’s eye view: some lessons from *Chlamydia psittaci*. FEBS Lett. 590, 3920–3940 (2016).

6. Elwell, C., Mirrashidi, K. & Engel, J. *Chlamydia* cell biology and pathogenesis. Nat. Rev. Microbiol. 14, 385–400 (2016).

7. Moore, E. R. & Ouellette, S. P. Reconceptualizing the chlamydial inclusion as a pathogen-specified parasitic organelle: An expanded role for Inc proteins. Front. Cell. Infect. Microbiol. 4, 1–10 (2014).

8. Heuer, D., et al. *Chlamydia* causes fragmentation of the Golgi compartment to ensure reproduction. Nature 457, 731–735 (2009).

9. Abdelrahman, Y., Ouellette, S. P., Belland, R. J. & Cox, J. V. Polarized Cell Division of *Chlamydia trachomatis*. PLoS Pathog. 12, e1005822 (2016).

10. Christian, J. et al. Targeting of a Chlamydial Protease Impedes Intracellular Bacterial Growth. PLoS Pathog. 7, e1002283 (2011).

11. Heymann, J., Lipinski, A., Bauer, B., Meyer, T. & Heuer, D. *Chlamydia trachomatis* infection prevents front-rear polarity of migrating HeLa cells. Cell. Microbiol. 15, (2013).

12. Rejman Lipinski, A., et al. Rab6 and Rab11 Regulate *Chlamydia trachomatis* Development and Golgin-84-Dependent Golgi Fragmentation. PLOS Pathog. 5, e1000615 (2009).

13. Koch-Edelmann, S. et al. The cellular ceramide transport protein CERT promotes *Chlamydia psittaci* infection and controls bacterial sphingolipid uptake. Cell. Microbiol. 19, 1–13 (2017).

14. Scholz, J., Holland, G., Laue, M., Banhart, S. & Heuer, D. *Chlamydia*-containing spheres are a novel and predominant form of egress by the pathogen *Chlamydia psittaci*. mBio 15, e01288–24 (2024).

15. Hybiske, K. & Stephens, R. S. Mechanisms of host cell exit by the intracellular bacterium *Chlamydia*. Proc. Natl. Acad. Sci. U. S. A. 104, 11430–11435 (2007).

16. Lutter, E. I., Barger, A. C., Nair, V. & Hackstadt, T. *Chlamydia trachomatis* Inclusion Membrane Protein CT228 Recruits Elements of the Myosin Phosphatase Pathway to Regulate Release Mechanisms. Cell Rep. 3, 1921–1931 (2013).

17. Nguyen, P. H., Lutter, E. I. & Hackstadt, T. *Chlamydia trachomatis* inclusion membrane protein MrcA interacts with the inositol 1,4,5-trisphosphate receptor type 3 (ITPR3) to regulate extrusion formation. PLoS Pathog. 14, 1–19 (2018).

18. Zuck, M., Sherrid, A., Suchland, R., Ellis, T. & Hybiske, K. Conservation of extrusion as an exit mechanism for *Chlamydia*. Pathog. Dis. 74, 1–4 (2016).

19. Derré, I. *Chlamydiae* interaction with the endoplasmic reticulum: contact, function and consequences. Cell. Microbiol. 17, 959–966 (2015).

20. Derré, I., Swiss, R. & Agaisse, H. The lipid transfer protein CERT interacts with the *Chlamydia* inclusion protein IncD and participates to ER-Chlamydia inclusion membrane contact sites. PLoS Pathog. 7, e1002092 (2011).

21. Agaisse, H. & Derré, I. Expression of the effector protein IncD in *Chlamydia trachomatis* mediates recruitment of the lipid transfer protein CERT and the endoplasmic reticulum-resident protein VAPB to the inclusion membrane. Infect. Immun. 82, 2037–2047 (2014).

22. Murray, R., Flora, E., Bayne, C. & Derré, I. IncV, a FFAT motif-containing *Chlamydia* protein, tethers the endoplasmic reticulum to the pathogen-containing vacuole. Proc. Natl. Acad. Sci. U. S. A. 114, 12039–12044 (2017).

23. Gudlur, A. et al. Calcium sensing by the STIM1 ER-luminal domain. Nat. Commun. 9, 4536 (2018).

24. Stiber, J. et al. STIM1 signalling controls store-operated calcium entry required for development and contractile function in skeletal muscle. Nat. Cell Biol. 10, 688–697 (2008).

25. Agaisse, H. & Derré, I. STIM1 is a novel component of ER-*Chlamydia trachomatis* inclusion membrane contact sites. PLoS One 10, 1–18 (2015).

26. Chamberlain, N. B., Dimond, Z. & Hackstadt, T. *Chlamydia trachomatis* suppresses host cell store-operated Ca^2+^ entry and inhibits NFAT/calcineurin signaling. Sci. Rep. 12, 1–11 (2022).

27. Cortina, M. E., Clayton Bishop, R., DeVasure, B. A., Coppens, I. & Derre, I. The inclusion membrane protein IncS is critical for initiation of the *Chlamydia* intracellular developmental cycle. PLoS Pathog. 18, 1–20 (2022).

28. Cortina, M. E. & Derré, I. Homologues of the *Chlamydia trachomatis* and *Chlamydia muridarum* Inclusion Membrane Protein IncS Are Interchangeable for Early Development but Not for Inclusion Stability in the Late Developmental Cycle. mSphere 8, (2023).

29. Elwell, C. A., et al. *Chlamydia trachomatis* Co-opts GBF1 and CERT to Acquire Host Sphingomyelin for Distinct Roles during Intracellular Development. PLOS Pathog. 7, e1002198 (2011).

30. Rockey, D. D., Fischer, E. R. & Hackstadt, T. Temporal analysis of the developing *Chlamydia psittaci* inclusion by use of fluorescence and electron microscopy. Infect. Immun. 64, 4269– 4278 (1996).

31. Hybiske, K. & Stephens, R. S. Exit strategies of intracellular pathogens. Nat. Rev. Microbiol. 6, 99–110 (2008).

32. Kumar, Y. & Valdivia, R. H. Leading a sheltered life: intracellular pathogens and maintenance of vacuolar compartments. Cell Host Microbe 5, 593–601 (2009).

33. Petit, T. J. P. & Lebreton, A. Adaptations of intracellular bacteria to vacuolar or cytosolic niches. Trends Microbiol. 30, 736–748 (2022).

34. Kumagai, K., Elwell, C. A., Ando, S., Engel, J. N. & Hanada, K. Both the N- and C- terminal regions of the Chlamydial inclusion protein D (IncD) are required for interaction with the pleckstrin homology domain of the ceramide transport protein CERT. Biochem. Biophys. Res. Commun. 505, 1070–1076 (2018).

35. Kumagai, K. et al. Chlamydial Infection-Dependent Synthesis of Sphingomyelin as a Novel Anti-Chlamydial Target of Ceramide Mimetic Compounds. Int. J. Mol. Sci. 23, (2022).

36. Banhart, S. et al. Improved plaque assay identifies a novel anti-*Chlamydia* ceramide derivative with altered intracellular localization. Antimicrob. Agents Chemother. 58, 5537–5546 (2014).

37. Goellner, S. et al. Transcriptional response patterns of *Chlamydophila psittaci* in different in vitro models of persistent infection. Infect. Immun. 74, 4801–4808 (2006).

38. Lienard, J. et al. Development of a new *Chlamydiales*-specific real-time PCR and its application to respiratory clinical samples. J. Clin. Microbiol. 49, 2637–2642 (2011).

39. Ran, F. A. et al. Genome engineering using the CRISPR-Cas9 system. Nat. Protoc. 8, 2281–2308 (2013).

40. Niekamp, P. et al. Ca^2+^-activated sphingomyelin scrambling and turnover mediate ESCRT-independent lysosomal repair. Nat. Commun. 13, (2022).

41. Scharte, F., Franzkoch, R. & Hensel, M. Flagella-mediated cytosolic motility of *Salmonella enterica* Paratyphi A aids in evasion of xenophagy but does not impact egress from host cells. Mol. Microbiol. 1–18 (2023) doi:10.1111/mmi.15104.

